# Integrated analysis of the miRNA-mRNA network associated with LMP1 gene in nasopharyngeal carcinoma

**DOI:** 10.1101/615237

**Authors:** Yang Yang, Wen Liu, Yan Zhang, Shuo Wu, Bing Luo

## Abstract

Epstein-Barr virus oncogenic latent membrane protein 1 (LMP1) has been known to play important roles in nasopharyngeal carcinoma (NPC). LMP1 gene also induced a variety of microRNAs (miRNAs) which bear pivotal roles in regulation of mRNAs expression. However, little was known about the global change of mRNAs and miRNAs induced by LMP1 gene in NPC. In our study, one NPC tissue microarray profile and two LMP1-associated microarray expression profiles data were downloaded from the Gene Expression Omnibus database. A protein-protein interaction network was constructed by using bioinformatics platform Gene-Cloud of Biotechnology Information (GCBI). 78 differentially expressed miRNAs and 3322 differentially expressed genes were identified in order to generate a macroscopic network between miRNAs and mRNAs associated with LMP1 gene. In addition, two significant models were generated to illustrate the expression tendency. Our study provided a way to reveal the interaction between miRNAs and mRNAs in LMP1 axis, bringing insights into the pathogenesis of NPC.

## Introduction

Nasopharyngeal carcinoma is a primary cancer arising from the nasopharynx. The disease commonly distributes in certain regions of East Asia and Africa, especially in Guangdong province of China [1]. It is generally considered that genetic susceptibility, Epstein-Barr virus (EBV) infection and environmental factors were involved in the pathogenesis of NPC. Linkage analysis showed that several susceptibility loci were related to the etiology of NPC in China [2,3]. Consumption of salt-preserved fish may also play an important role in NPC [4].

EBV is an enveloped, double-stranded human DNA herpesvirus. It infects more than 90% of the global adult people being for life and in most carriers the infection is usually asymptomatic. Nevertheless, EBV has oncogenic properties for several malignancies including Hodgkin’s disease, Burkitt’s lymphoma, as well as undifferentiated NPC [5]. The consistent detection of virus in NPC cases and the continuous expression of virus genes indicate that EBV is crucial for the malignant growth. The circulating cell free EBV DNA is well-recognized biomarker in undifferentiated NPC [6]. EBV latently infects NPC cells by expressing multiple virus genes including EBV nuclear antigens 1 (EBNA1), the latent membrane proteins (LMP1), 2A, and 2B and viral noncoding RNAs. Among them, LMP1 oncoprotein may play a critical role in the initiation and progression of NPC as well as the invasion and metastasis [7–9]. LMP1 is a 66kDa integral membrane protein consisting of a cytoplasmic amino-terminus, six transmembrane domains and a large cytoplasmic C-terminal tail comprising three regions: transformation effector site (TES)/C-terminal activating regions (CTAR) 1, 2 and 3. CTAR1, CTAR2 are responsible for recruiting cellular signaling molecules of the tumor necrosis factor receptor associated factor (TRAF) family and TRAF-associated death domain protein (TRADD) and the activation of the nuclear factor kap-pa-light-chain-enhancer of activated B cells (NF-κB) and c-Jun N-terminal protein kinase (JNK) /DNA-binding activity of activator protein-1 (AP-1) signaling [10,11].

It is well recognized that the many small non-coding microRNAs (miRNAs) play pivotal role in regulation the homeostasis of mRNAs expression in physiological and pathological process [12]. MiRNAs can bind to the specific regions in the target mRNAs, thus leading to the degradation or repression of the mRNAs [13]. As a multifunctional gene, LMP1 was proved to modulate many kinds of miRNAs in B cells and epithelial cells [14–16]. Most of these studies have focused on the interaction of single miRNA with the carcinoma cells, thus a landscape of interaction between miRNAs and mRNAs was needed to illustrate the global change induced by LMP1 in NPC.

In the present study, we investigated three expression profiles involving LMP1-associated miRNAs and mRNAs data and NPC tissue samples from the Gene Expression Omnibus (GEO) database (http://www.ncbi.nlm.nih.gov/geo/). We analyzed the differentially expressed genes (DEGs) and miRNAs (DE-miRNAs) and performed the pathway analysis. The protein-protein interaction (PPI) network and the mRNA-miRNA interaction network were constructed to obtain the key genes and miRNAs in NPC.

## Materials and Methods

### Data Collection

Gene expression profiles were downloaded as raw signals from GEO. The dataset GSE12452 comprised 31 NPC tissues and 10 normal nasopharyngeal tissues. The dataset GSE29297 involved a series of LMP1 TES2-stimulated HEK-293 cell line samples at different time points. The dataset GSE26596 valuated the miRNAs expression in NPC cell line TW03 transfected with LMP1. TW03 was an EBV-negative cell line derived from a lymphoepitheliomatous undifferentiated carcinoma in Taiwan province [17]. The sequences of the LMP1 gene encoded by different EBV strains has been shown to have a degree of variation in NPC. The LMP1 vector used in dataset GSE29596 was as described [18]. Vector encoding LMP1 TES2 (amino acids 351-386) was used in GSE29297. Both of the two vector covered part of C-terminal activating region.

### Differencial Expression Analysis

An online platform Gene-Cloud of Biotechnology Information (GCBI) was used to interpreted, normalized, log2 scaled the datasets GSE12452 and GSE29297. This platform integrated with biology, computer science, medicine, informatics, mathematics and other disciplines. Microarray probe signals with absolute value of fold change (FC)>1.2, *P*-value<0.05, and false discovery rate (FDR) <0.05 were considered to be statistically differential [19,20]. Co-expression networks were constructed based upon contribution degrees and models of Series Test of Cluster (STC) were established by random permutation scheme (http://college.gcbi.com.cn/helpme). GSE26596 was analysed via the R-programming language-based dataset analysis tool GEO2R (http://www.ncbi.nlm.nih.gov/geo/geo2r/). This interactive tool was allowed to screen different expression miRNAs between TW03 cell line and TW03 cell line transfected with LMP1 in dataset GSE26596 [21].

### GO and pathway analysis

The functional annotation of the DEGs and DE-miRNAs were performed by GCBI platform. Gene Ontology (GO) (http://www.geneontology.org) classification was used to analysis the DEGs function including biological processes, molecular function, and cellular component [22]. The Kyoto Encyclopedia of Genes and Genomes (KEGG) pathway enrichment analysis was utilized to explore the pathway of DEGs.

### Gene regulation network and miRNA-mRNA network

Micro RNA target genes were predicted mainly based on the bioinformatic platform miRWalk 2.0 [23]. The platform integrates information from 12 miRNA-target databases, including prediction datasets and validated information: MiRWalk, MicroT4, miRanda, miRBridge, miRDB, miRMap, miRNAMap, PICTAR2, PITA, RNA22, RNAhybrid, Targetscan. The 3’ untranslated regions for the target genes were used as the primary base-pairing regions of miRNAs [24]. The miRNA-gene pairs that were common in at least five databases and met with *P*<0.05 were regard as reliable. The miRNA target mRNAs were then compared with the genes in STC models and the overlapped genes were filtered for further analysis. The LMP1 associated miRNA-mRNA interactomes was further visualized using the Cytoscape 3.5.1 platform [25]. In addition, DEGs of GSE29297 and GSE12452 were used to construct a protein-protein interaction networks based on GCBI platform. The details of the network construction algorithm were described on GCBI platform.

## Results

### Identification of DEGs and DE-miRNAs associated with LMP1

In the present study, the mRNA profiling dataset GSE29297 was analyzed on GCBI platform. Samples in the dataset were divided into five groups according to the TES2 stimulation period. 3109 probes for 2787 DEGs were identified after background correction and normalization (*P* value<0.05, Q value<0.05, Figure 1). 41 samples (including 31 NPC samples and 10 normal tissues) in the dataset GSE12452 were divided into two groups using GCBI tools. 2566 probes for 1815 genes were up-regulated and 2021 probes for 1507 genes were down-regulated. The criteria were set to adjust *P* value <0.05 and FC>1.5 (Figure 2). 372 DEGs were up-regulated and 245 DEGs were down-regulated in both datasets GSE29297 and GSE12452. For dataset GSE26596, 72 miRNAs were up-regulated, and 6 miRNAs were down-regulated (adjust *P* value<0.05, FC>1.5).

**Figure 1.**
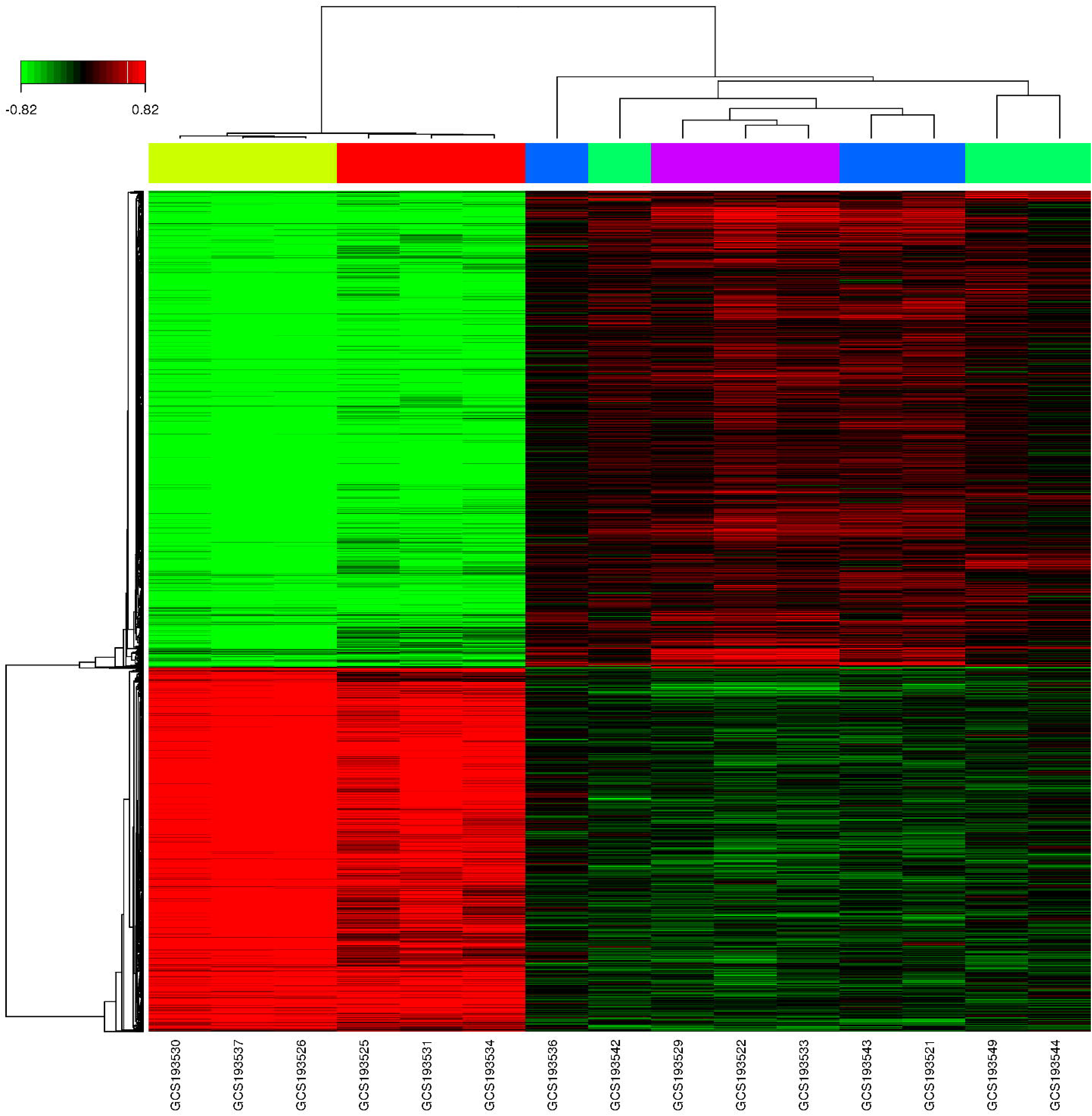
Heat map showing the differential expression pattern of genes in dataset GSE29297. The x-axis represents samples and the y-axis represents the genes. The bar on the top indicates the log2-scaled expression level, up-regulated genes were presented in red, while down-regulated in green.

**Figure 2.**
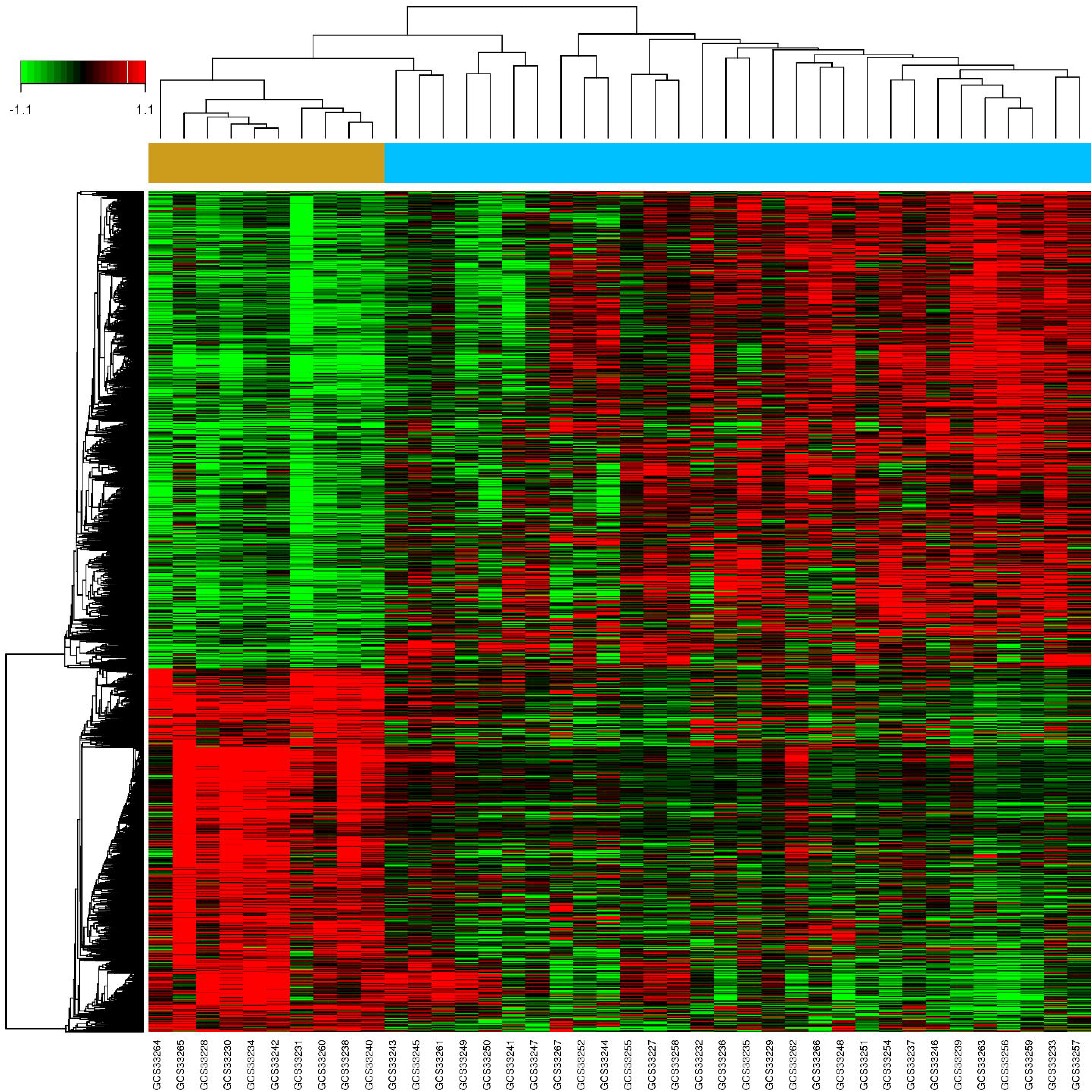
Heat map showing the differential expression pattern of genes in dataset GSE12452. The x-axis represents samples and the y-axis represents the genes. The bar on the top indicates the log2-scaled expression level, up-regulated genes were presented in red, while down-regulated in green.

### Models of Series Test of Cluster

The expression tendency of gene clusters of dataset GSE29297 was calculated on GCBI platform. Among the 20 expression STC models constructed, 15 models were estimated as statistical significant (all *P* values<0.05, Figure 3). Notably, a descending expression model STC32 including 1358 probes for 1070 genes and an ascending model STC49 including 1422 genes were identified as significantly enriched, which may be affected by the LMP1 associated up-regulated miRNAs.

**Figure 3.**
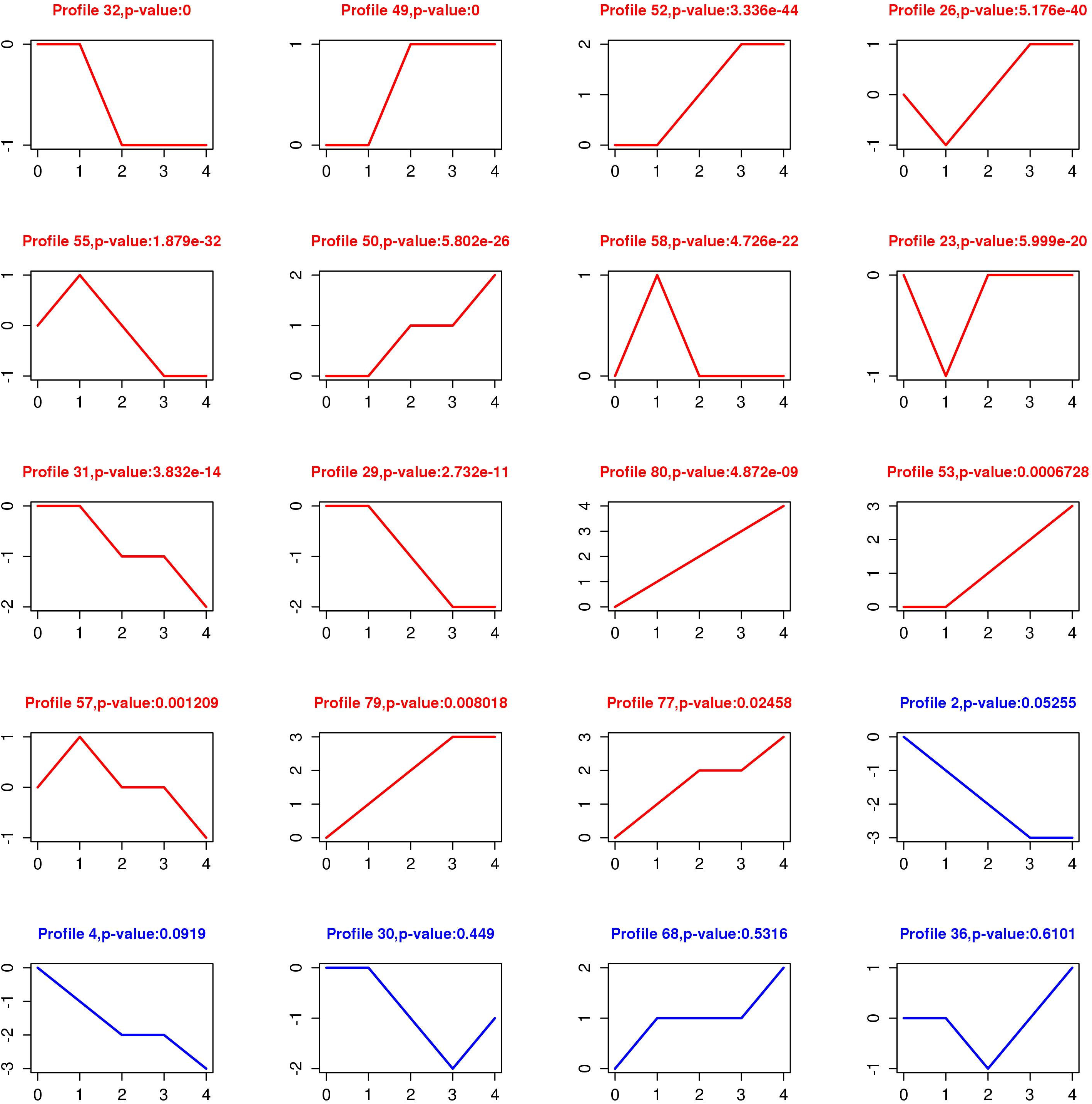
STC models. The profile represent the series test of cluster (STC), profile number represent the STC models. P-value: adjusted P-value represents the significant levels of the genes in a certain model compared with theoretical genes. The diagram was present in red lines as the P-value of the model<0.05, otherwise in blue lines. Numbers in horizontal ordinate were relative periods of the LMP1-TES2 gene stimulation on the HEK293 cell line. The ordinate numbers represent relative grades of expression levels.

### Prediction of miRNA-target genes

The potentially targeted mRNAs were predicted using the integrated bioinformatic tool miRWalk 2.0. Among the 78 DE-miRNAs, 11 miRNAs showed up-regulated expression with at least 20 fold change, including hsa-miR-134, hsa-miR-7, hsa-miR-432 and miR-515 family such as hsa-miR-516-3p, hsa-miR-520b, hsa-miR-520d, hsa-miR-520e, hsa-miR-523. While only 6 miRNAs down-regulated mildly, including the hsa-miR-15a, hsa-miR-107, mmu-miR-140*, hsa-miR-149, hsa-miR-194, hsa-miR-103. The target genes of several miRNAs were later observed with the STC models.

### Significant functions and pathway enrichment analysis

Gene ontology and KEGG pathway enrichment analysis were performed on the DEGs to reveal their biological significance in NPC on GCBI platform.

The results showed that the DEGs in dataset GSE29297 were significantly involved in the molecular functions of protein binding, zinc ion binding and DNA binding and sequence-specific DNA binding transcription factor activity. In terms of biological processes, the DEGs were mainly enriched in the process of positive and negative regulation of transcription. Mitogen-activated protein kinase (MAPK) signaling pathway, phosphatidylinositol-3-kinase-Serine/threoninekinase (PI3K-AKT) signaling pathway and pathways of human T-cell leukemia virus type 1 (HTLV-I) and hepatitis B were predominately significant in the KEGG pathway analysis.

GO enrichment analysis revealed that the DEGs of dataset GSE12452 were significantly enriched in protein and ATP binding, DNA and RNA binding as well as mitotic cell cycle. KEGG pathway analysis showed that the DEGs were significantly enriched in the metabolic pathway, the PI3K signaling pathway and pathways associated with cell cycle and RNA transport.

### Network Analysis

The protein-protein coexpression network showed that a number of genes presented higher node degree including exportin 1(XPO1), eukaryotic translation initiation factor 3(EIF3E), transcription elongation regulator 1(TCERG1), anaphase promoting complex subunit 5 (ANAPC5), cytochrome c, somatic (CYCS) (Figure 4). The pathway relation network showed that MAPK signaling pathway, pathway of cell cycle and apoptosis and p53 signaling pathway were significant-enriched downstream pathways, while the pathway in cancer regulated significantly upstream (Figure 5).

**Figure 4.**
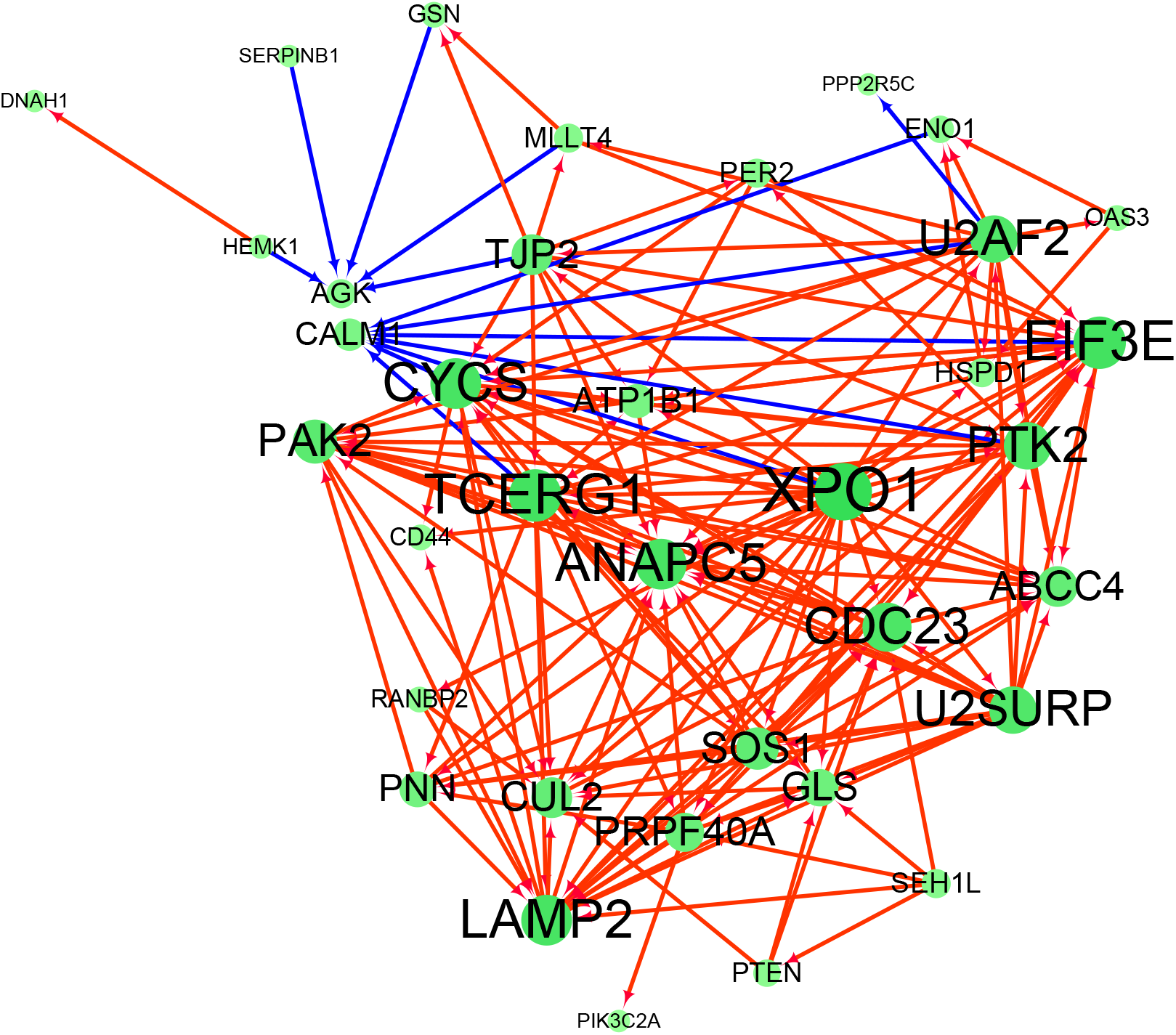
Gene co-expression network of datasets GSE29297 and GSE12452.The size of the node represents the degree of the gene. The positive correlation coefficient was presented in red line, the negative was in blue.

**Figure 5.**
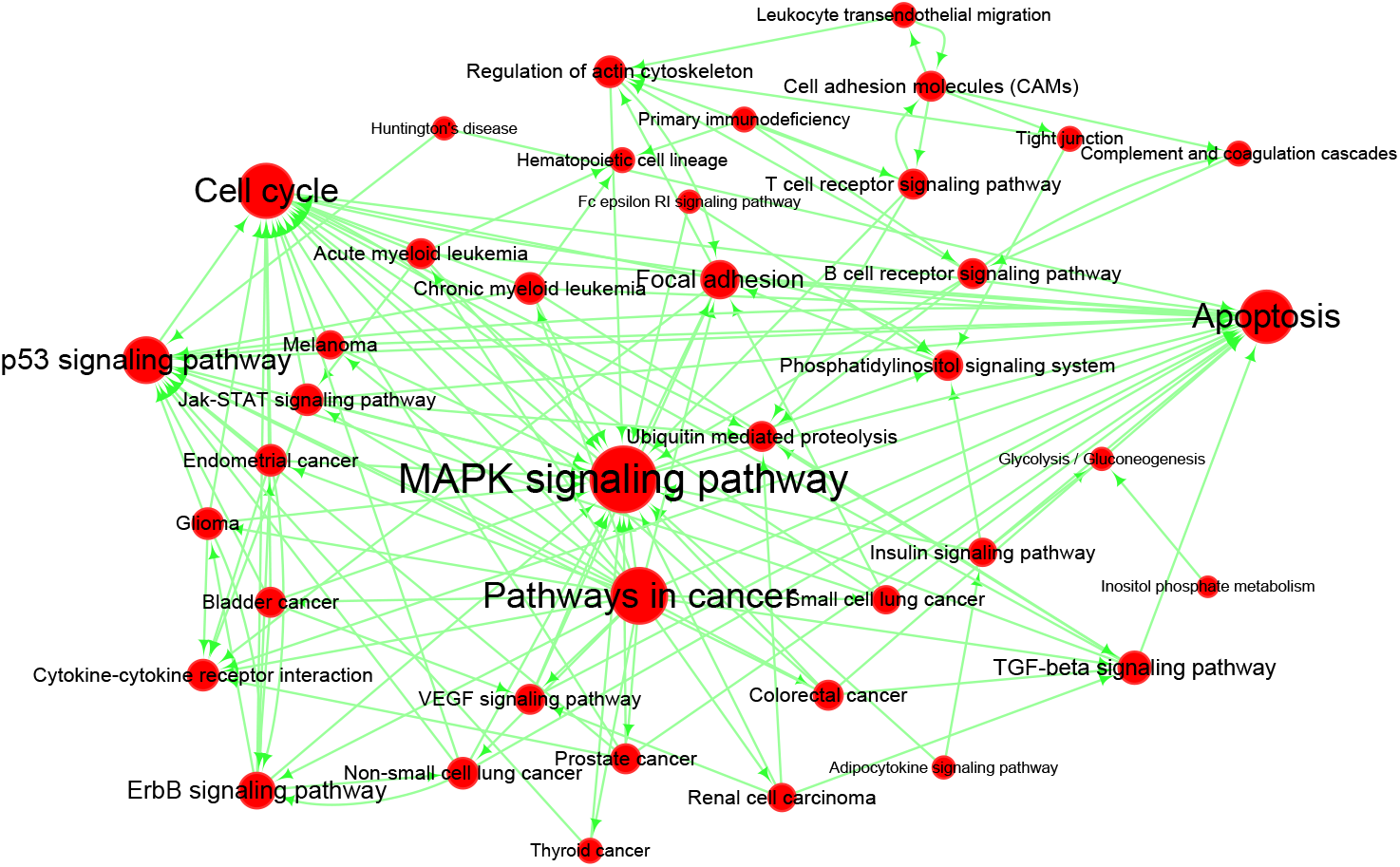
Pathway relation network of datasets GSE29297 and GSE12452. The arrow in the network directed to the downstream pathways. The size of the node represents the degree of pathway.

The time-associated DEGs in STC32 and STC49 models and the up-regulated miRNAs identified in GSE26596 were used to generate a LMP1-specific miRNA-RNA interacting network. Top 5 miRNAs were chosen as core miRNAs to built the network (Figure 6).

**Figure 6.**
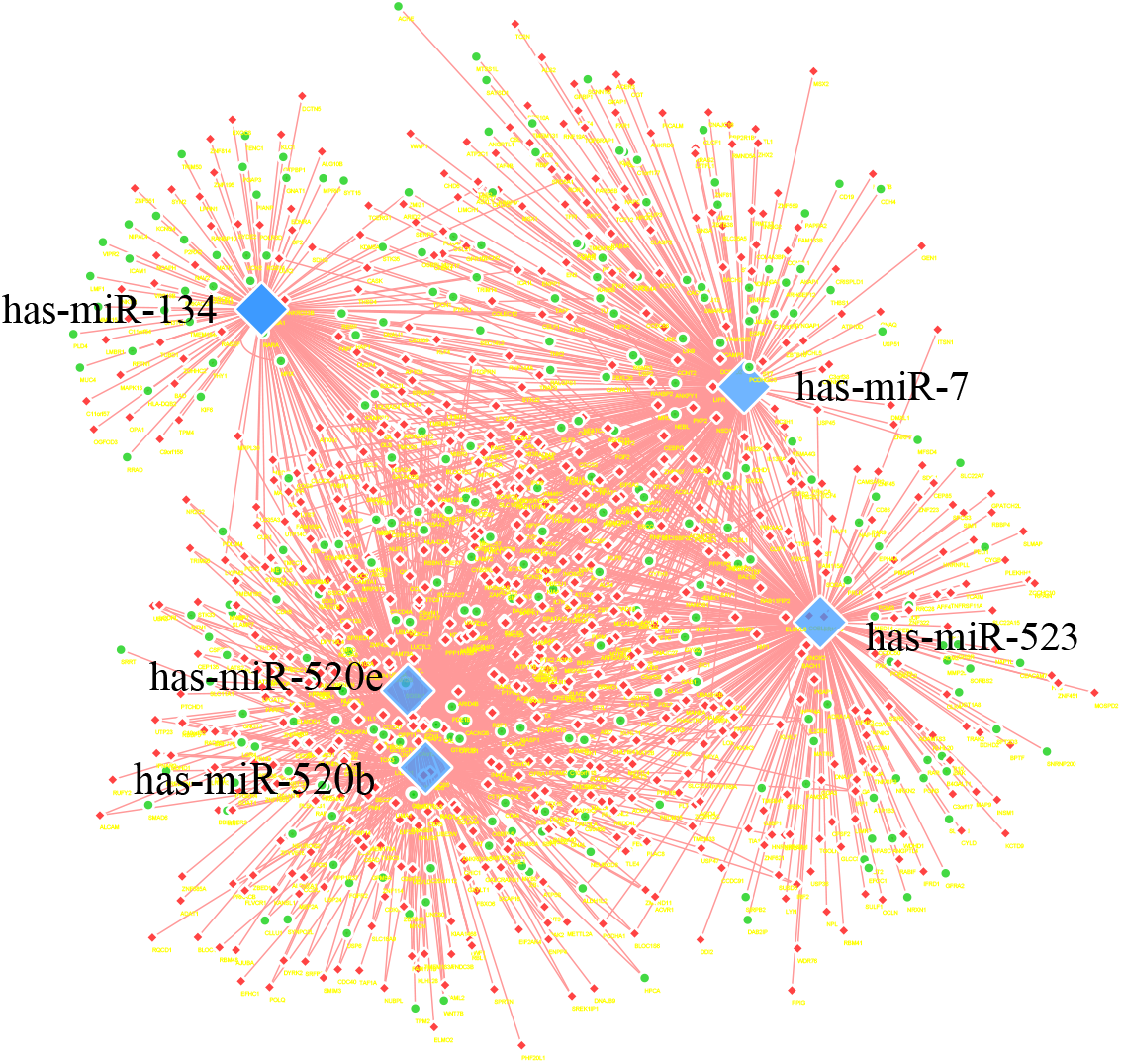
Interaction network between top 5 up-regulated miRNAs in GSE26596 with expression models STC32 and STC49. Up-regulated genes were filled in Red diamond, and down-regulated genes green ellipse. The miRNAs were filled in blue diamond. These genes were regarded as targets of the miRNAs in predicted and validate databases as described.

## Discussion

NPC featured an EBV type □ latency which was characterized by the expression of LMP1 [7]. High level miRNAs such as the BamHI-A rightward transcripts (BARTs), with the combination of the human noncoding RNAs induced by the virus, may affect the process of the oncogenesis. However, a variety of ncRNAs may hinder our understanding about the LMP1 axis. This integrated study aimed to seek a clue about the interaction between LMP1 and its induced miRNAs in the universe of the signal network.

A co-expression analysis of the datasets GSE12452 and GSE29297 displayed a 36-hub genes network. Among these hub genes, the major export receptors of mRNA (XPO1) gene was predicted to co-express with 22 genes in NPC cell and the expression level was 1.8 fold up-regulated (*P*<0.01). The gene can be regulated by hsa-miR-155 and hsa-miR-373*, which were upregulated in GSE26596. This dataset comprised the expression profile of EBV-negative NPC cell line TW03 and the LMP1-transfected TW03 cell line. We proposed that the up-regulated expression levels of miRNAs act as a response to attenuate the up regulated mRNAs such as the XPO1.

An overlapped analysis of the datasets GSE12452 and GSE29297 revealed that the DEGs were significantly involved in the PI3K-AKT signaling pathway. Studies have shown that LMP1 can activate the PI3K-AKT pathway and evade tumor suppressor responses in many ways. LMP1 over-expression led to upregulated level of mi-155-Ubiquilin1 axis to activate PI3K-AKT pathway in NPC cells, which promoted the radioresistence of NPC [26]. In diffuse large B-cell lymphoma the PI3K-AKT signaling pathway was significantly activated by the overexpression of miR-155 in DHL16 cells, while in OCL-Ly3 cells knockdown of miR-155 attenuate the AKT activity [27]. Correspondingly, in GSE26596, miRNA-155 was 12.73 fold upregulated (adjust *P*<0.01), which indicating that the PI3K-AKT signaling pathway may possibly be regulated by LMP1 via miR-155 in NPC.

Analysis showed that most genes in GSE29297 were enriched in two STC expression models in LMP1 TES2-stimulated HEK-293 cell line, named STC32 and STC49. The ascendant expression model STC49 indicated that 1787 probes for 1422 genes were up-regulated in pace with the TES2 stimulation. Pathway analysis showed that these genes were primarily enriched in the processes such as ubiquitin mediated proteolysis and transforming growth factor-signaling pathway. It has been proved that LMP1 can regulate ubiquitination by the activation of Interferon regulatory factor 7 [28]. While in the down-regulated gene model STC32, the genes were enriched in the process of mesenchymal to epithelial transition and cell communication. Several studies have shown that LMP1 can promote the progress of epithelial to mesenchymal transition, thus contributes to the metastatic nature of NPC [29,30].

Integrate analysis revealed a miR-515 family including hsa-miR-520e, hsa-miR-523, hsa-miR-520b, hsa-miR-516-3p and hsa-miR-520d was significantly up-regulated. The targets of miR-515 family target genes were predicted to cover large number of genes in the down-regulated model STC32. However, as shown in figure.4, more putative interactions were in the ascending model STC49, hinting that the up-regulation of miRNAs may be a restrain means or a remedial measure to counteract the up-regulation of particular mRNAs. For example, analysis of miRNA hsa-miR-520e targeted genes showed that the down-regulated genes were primarily enriched in glypican pathway, while the up-regulated genes were mainly enriched in vascular endothelial growth factor signaling pathway, hinting that hsa-miR-520e may play a negative role in balancing the abnormal increasement of specific miRNAs. Another study showed that miR-520 acted as a tumor-suppressive factor by direct targeting of transcription factor P65 and thus inhibit the NF-κB signaling to reduce the expression of the pro-inflammatory cytokines [31].

Recently, it has been reported that hsa-miR-134 played crucial roles in abundant and complicated pathways, including KRAS signal pathway, Notch pathway and EGFR pathway [32,33]. In dataset GSE2927, both up-regulated genes such as SMAD6, MYCN, PRLR, CYTH3, BMP3, PDE5A, and down-regulated genes such as BCL2, TGFB2, FOXP2, PKD2, TAF4B, and TSN1 were predicted to be targets of miR-134. As many studies showed, miR-134 not only functions as a tumor repressor, but also acts as a cancer promoter. High expression of miR-134 contributes to head and neck carcinogenesis by targeting the WW Domain-Containing Oxidoreductase (WWOX) gene [34]. However, overexpression of miR-134 can inhibit the cell cycle progression of human human ovarian cancer stem cells and decrease the tumorigenicity in nude mice [35]. These studies hinting that a disregulation of miR-134 may participate in the development of NPC.

Nevertheless, our study had its points which were not rigorous enough, for example, the dataset GSE29297 used was limited in only one cite TES2, the DEGs and the DE-miRNAs involved in our study might associate with the sample sorts, the pathological stage and the condition of cell lines. In addition, more experiments were necessary to validate our results.

## Conclusion

In conclusion, by taking advantage of bioinformatic tools and GEO profiles, a protein-protein network and a miRNA-mRNA interaction network were constructed. Our result revealed another layer of gene regulation network in the LMP1-associated gene expression axis, which would provide a better understanding of the interaction of mRNAs and miRNAs. The interaction relationship may open a way to explore the potential use of miRNA in NPC.

## Acknowledgments

This research was supported by the National Natural Science Foundation of China (NSFC 81571995) and the Natural Science Foundation of Shandong Province (ZR2015HM069). The authors appreciated Linlin Wang for providing an account of GCBI tools.

## Authors’ contributions

BL and QS conceived the project and designed experiments. WL, YZ, SW and YY participated in performance of the experiments and the drafting of the manuscript. All authors read and approved the final manuscript.

## Ethics, consent and permissions

### Ethical approval

The study was approved by Ethics Committee of Qingdao University Medical College.

### Consent to publish

All the datasets used in the study were obtained from GEO public database, which were equipped with informed consent.

